# Cellular and transcriptional profiles of peripheral blood mononuclear cells pre-vaccination predict immune response to preventative MUC1 vaccine

**DOI:** 10.1101/2024.06.14.598031

**Authors:** Daniel Y. Yuan, Michelle L. McKeague, Vineet K. Raghu, Robert E. Schoen, Olivera J. Finn, Panayiotis V. Benos

## Abstract

A single arm trial (NCT007773097) and a double-blind, placebo controlled randomized trial (NCT02134925) were conducted in individuals with a history of advanced colonic adenoma to test the safety and immunogenicity of the MUC1 tumor antigen vaccine and its potential to prevent new adenomas. These were the first two trials of a non-viral cancer vaccine administered in the absence of cancer. The vaccine was safe and strongly immunogenic in 43% (NCT007773097) and 25% (NCT02134925) of participants. The lack of response in a significant number of participants suggested, for the first time, that even in a premalignant setting, the immune system may have already been exposed to some level of suppression previously reported only in cancer. Analysis of single-cell RNA-sequencing (scRNA-seq) data from banked pre-vaccination peripheral blood mononuclear cells (PBMCs) (16 immune responders and 16 non-responders) identified specific cell types, genes, and pathways of a productive vaccine response. Responders had a significantly higher percentage of CD4+ naïve T cells pre-vaccination, but a significantly lower percentage of CD8+ T effector memory (TEM) cells and CD16+ monocytes. Differential gene expression (DGE) and transcription factor inference analysis showed a higher level of expression of T cell activation genes, such as Fos and Jun, in CD4+ naïve T cells. Pathway analysis showed enriched signaling activity in responders. Furthermore, Bayesian network analyses suggested that these genes were mechanistically related to response. Our analyses identified several immune mechanisms and candidate biomarkers which can be further validated as predictors of immune responses to a preventative cancer vaccine that could facilitate selection of individuals likely to benefit from a vaccine or be used in further research to improve vaccine responses.

**One Sentence Summary:** Single-cell RNA sequencing reveals distinctive cell types, enriched biological pathways, and candidate biomarkers pre-vaccination that predict immune responses to the preventative MUC1 cancer vaccine.

## INTRODUCTION

Cancer immunotherapy is a rapidly evolving field of research that is leading to prompt clinical applications. In 2022-2023, there were over 14 new approved immunotherapies (*1*). While these treatments have led to cures and long-term remissions in some patients in some cancers (*2*), the majority of patients do not experience therapeutic benefit (*3, 4*). The main reason for the lack of therapeutic efficacy is the reduced ability of the immune system of cancer patients to fight cancers that have already escaped immune control (*5*). Chronic exposure of the immune system to cancer, combined with the highly immunosuppressive tumor microenvironment, leads to exhaustion or elimination of cancer specific effector T and B cells and accumulation of regulatory T cells (Treg), myeloid-derived suppressor cells (MDSCs), and immunosuppressive tumor promoting cytokines (*6*).

Current immunotherapies are designed to reactivate cancer specific immunity by using check point inhibitors (*2*) and cytokine therapy (*7*) or by delivering, through adoptive transfer, cancer specific T cells (*8*), NK cells, (*9*) or antibodies (*10*). Most of these strategies are accompanied by serious side effects and high costs that for now limit their applicability (*11, 12*). Thus, despite these new conceptual and technological developments, cancer remains a major health problem world-wide. According to the predictions by the World Health Organization’s (WHO) International Agency for Research on Cancer (IARC), the incidence of cancer world-wide will increase in both males and females from close to 20 million in 2020 to close to 30 million in 2040 (*13*). Therefore, further improvements in current therapies and development of new immune-based strategies are needed.

The discovery of tumor antigens that could be targets of human T cells and antibodies (*14, 15*) stimulated ongoing research into cancer vaccines (*16*). Based on encouraging animal experiments, hundreds of phase I and II clinical trials were conducted, with some progressing to phase III; however, none showed good clinical efficacy and only one reached the approval stage (*17*). As was the case with many newer immunotherapies, it was determined that the immunosuppressive environment established in advanced cancer also compromised the success of therapeutic vaccines.

Unlike other forms of immunotherapy that require the presence of cancer, cancer vaccines could be reimagined for cancer prevention (*12*), similar to existing vaccines against pathogens and vaccines for the prevention of viral cancers, such as Human Papilloma Virus (HPV) and Epstein-Barr Virus (HBV). Based on antigens expected to be expressed only in specific cancers, such vaccines could generate cancer specific immune memory in individuals at high risk for those cancers before the malignant transformation and the accompanied immune suppression could take place. Many cancers are preceded by well-characterized premalignant changes that express cancer antigens, and vaccines given in that setting could elicit immune responses capable of eliminating premalignant lesions thus intercepting their progression to cancer (*18–20*).

Vaccines for prevention or interception of non-viral cancers are beginning to be tested in animal models and in early clinical trials. The first two clinical trials of a cancer vaccine applied in the absence of cancer, tested a peptide vaccine based on the MUC1 tumor associated antigen (*21, 22*). MUC1 vaccines had been previously tested as therapeutic vaccines in multiple Phase I and II clinical trials in advanced cancer patients, meeting safety criteria but without showing significant immunogenicity or clinical efficacy (*23–31*).

The first trial in a preventative setting was a single arm trial (NCT007773097) conducted from 2008 to 2012 (*21*) and the second was a multi-center double-blind, placebo controlled randomized trial (NCT02134925) conducted from 2015 to 2020 (*22*). In both trials the vaccine was administered to individuals with a recent history of advanced colonic adenoma, a premalignant precursor to colon cancer. The vaccine was safe and immunogenic, as measured by induction of anti-MUC1 IgG. The response rates of 43% in the first trial and 25% in the second trial were much higher than previously observed in cancer patients and the antibody titers were many folds higher, suggesting a much less suppressed immune microenvironment. Failure of this apparently immunogenic vaccine to elicit immune responses in other trial participants, suggested that some type of an immunosuppressive environment might already be present in the premalignant setting in those individuals. Extensive cellular analyses of the PBMC collected in both trials showed statistically significantly higher circulating levels MDSCs in non-responders as one important factor controlling the vaccine immunogenicity (7, 8).

In an initial exploratory study to understand additional differences between responders and non-responders (*32*), flow cytometry and bulk RNA sequencing data were obtained on banked PBMCs from both trials. The study showed that gene expression features from PBMCs 2 weeks post-vaccination had predictive value for the 12-week IgG response (AUROC=0.741) while bulk RNA-seq data on pre-vaccination samples was not predictive.

Seeking more sensitive methodology that would allow us to predict pre-vaccination who would respond to the vaccine, as well as elucidate the differences between responders and non-responders, we applied scRNA-seq on pre-vaccination PBMCs from 16 responders and 16 non-responders combined from both trials. Using standard bioinformatic approaches, such as differential gene expression and pathway analysis, we determined that, on average, vaccine responders had 34.7% more naïve CD4+ T cells, while non-responders had 87.9% more effector memory CD8+ T cells. Responders also demonstrated higher T cell activation, translational and transcriptional activity, and immune cytokine signaling. Graphical modeling supported these findings and identified several genes that directly affected response and could be used to build machine learning models to predict immune response at week 12 post-vaccination from pre-vaccination data.

## RESULTS

### scRNA-seq of pre-vaccination PBMC reveals immune cell population differences between vaccine responders and non-responders

We performed scRNA-seq on stored PBMC samples collected from patients right before the first vaccine injection (week 0). Two additional injections followed at week 2 and week 10. Response in both trials was defined by having at least a two-fold increase in anti-MUC1 IgG levels at week 12, compared to pre-vaccination (baseline). Clinicopathological data of the selected individuals are presented in **Table 1**. Overall quality of the sequencing can be seen in Figure S1.

**Table 1:** Patient Demographics

|  | **NR**<br>**(N=16)** | **R**<br>**(N=16)** |
| --- | --- | --- |
| **Trial** |  |  |
| NCT007773097 | 4 (25.0%) | 4 (25.0%) |
| NCT02134925 | 12 (75.0%) | 12 (75.0%) |
| **Age** |  |  |
| Mean (SD) | 60.4 (9.11) | 55.2 (6.98) |
| Median [Min, Max] | 65.0 [40.0, 70.0] | 53.5 [42.0, 67.0] |
| **Gender** |  |  |
| Female | 5 (31.3%) | 10 (62.5%) |
| Male | 11 (68.8%) | 6 (37.5%) |
| **BMI** |  |  |
| Mean (SD) | 28.6 (4.53) | 29.3 (4.44) |
| Median [Min, Max] | 29.0 [22.7, 36.5] | 29.1 [19.1, 37.2] |
| Normal | 5 (31.3%) | 2 (12.5%) |
| Obese | 5 (31.3%) | 7 (43.8%) |
| Overweight | 6 (37.5%) | 7 (43.8%) |
| **Ethnicity** |  |  |
| Hispanic | 3 (18.8%) | 3 (18.8%) |
| Non-Hispanic | 9 (56.3%) | 9 (56.3%) |
| Unknown or no information | 4 (25.0%) | 4 (25.0%) |
| **Race** |  |  |
| Black or African American | 1 (6.3%) | 0 (0%) |
| White | 11 (68.8%) | 11 (68.8%) |
| Unknown or no information | 4 (25.0%) | 5 (31.3%) |
| **IgG at week 0 (baseline)** |  |  |
| Mean (SD) | 0.218 (0.149) | 0.288 (0.144) |
| Median [Min, Max] | 0.209 [0.0400, 0.567] | 0.297 [0.0370, 0.634] |
| **IgG at week 12** |  |  |
| Mean (SD) | 0.211 (0.179) | 1.78 (1.27) |
| Median [Min, Max] | 0.131 [0.000444, 0.545] | 1.04 [0.326, 3.75] |
| **Ratio (IgG at week 12 vs week 0)** |  |  |
| Mean (SD) | 1.03 (0.357) | 7.55 (4.16) |
| Median [Min, Max] | 1.02 [0.258, 1.77] | 7.09 [1.65, 14.1] |

Principal component analysis (PCA) and uniform manifold approximation and projection (UMAP) dimensionality reduction visualization of the entire population without any correction demonstrated a significant batch variation as seen Figure S2. To maximize biological variation while mitigating technical variation, several batch integration methods were evaluated, which can be seen in Figure S3. The selected batch correction method and visualization results can be seen in **Figure 1A** and **Figure 1B**.

**Fig. 1:**
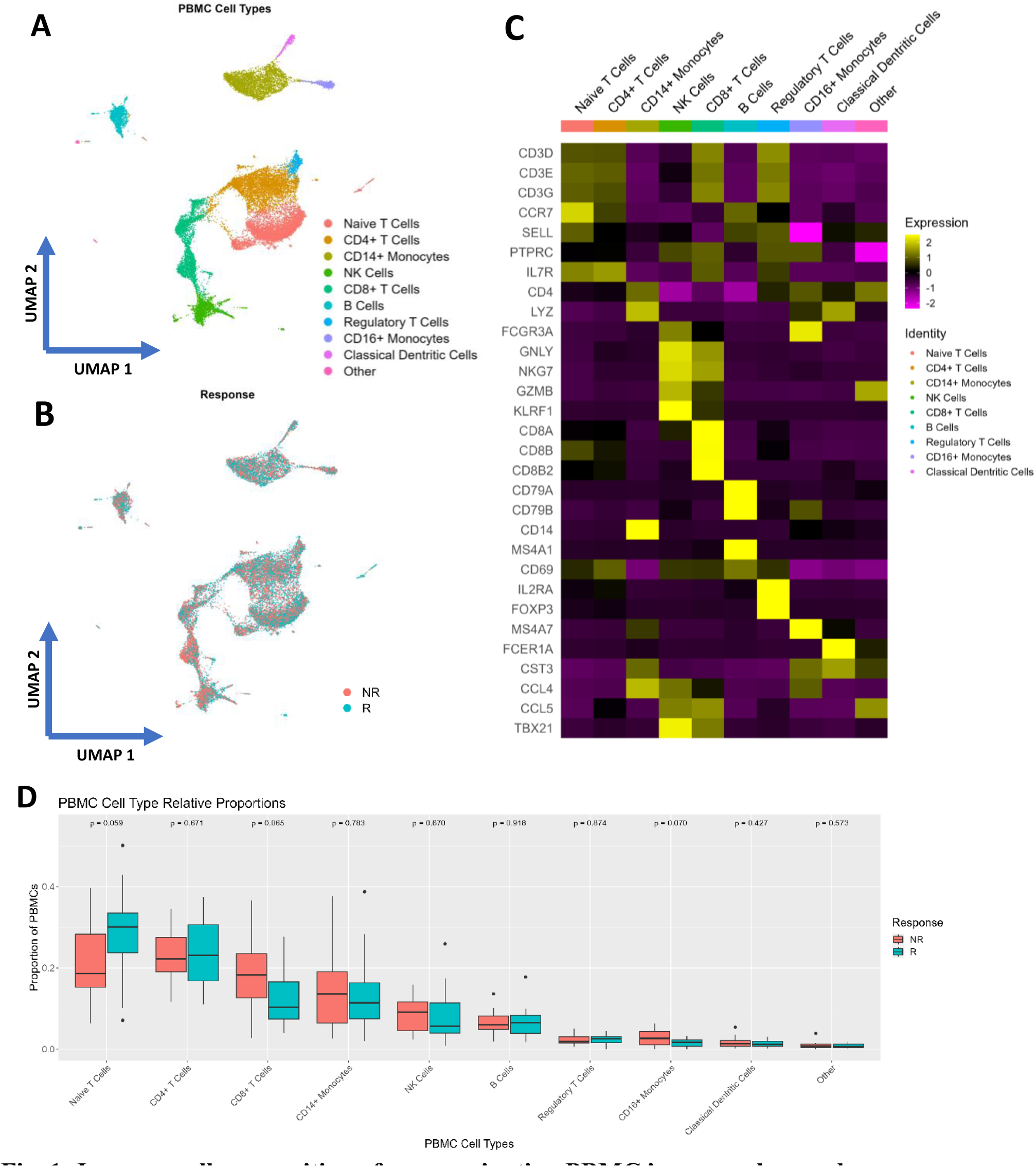
Immune cell composition of pre-vaccination PBMC in responders and non-responders. (**A** and **B**) Uniform manifold approximation and projection (UMAP) integrated embedding of 38,244 cells from PBMCs, colored by (A) cell type and (B) vaccine response. **(C)** Marker gene signatures in PBMC cell types (normalized averaged expression by cell type) **(D)** Paired responder vs non-responder box plot representing relative average proportion of each PBMC cell population, with p value calculated using t test

We ran shared nearest neighbor (SNN) based clustering with manual parameter selection by using several different resolution values and selecting the most stable resolution value for downstream tasks. The cluster membership stability across different resolution values can be seen in Figure S4A. After examining the clustering tree, we selected resolution 0.4 (see clusters in Figure S4B).

Subsequently, we determined the cell types using both manual and automated annotation. The markers for manual annotation can be seen in **Figure 1C**. The automated annotation mapped our dataset onto the existing multi-modal dataset (*33*) which can be seen in Figure S4C. The major groups identified correspond to the major cell types found in PBMCs – T and NK cells, B cells, and monocytic cells.

The labeled cell types were used to calculate the relative proportion of specific cell types in responders and non-responders, shown in **Figure 1D**. The manual annotation of the whole PBMC dataset allowed us to identify each major cell population, which was then subset and further clustered into individual subtypes. For example, within the monocyte subtypes, the CD16+ monocyte population is 64% higher in non-responders than responders with a slightly relaxed significance value (p = 0.07).

We repeated the same clustering process only on the T cell group. The manually annotated clusters and markers used are shown in **Figure 2A** and **Figure 2B**. To further distinguish between the different T cell subtypes, we used pseudotemporal analysis to calculate the estimated developmental trajectory of the overall T cell population, which can be seen in **Figure 2C** and Figure S5A. From the estimated trajectory, the CD8+ T effector memory (TEM) group 2 is a later stage of development of CD8+ TEM cells and corresponds to exhausted phenotypes. We also calculated differentially expressed genes (DEGs) for each individual T cell subtype and showed that the two CD8+ TEM populations were additionally characterized by their granzyme expression, which can be seen in Figure S5B. CD8+ TEM group 1 was GZMK+ while CD8+ TEM group 2 was GZMB+.

**Fig. 2:**
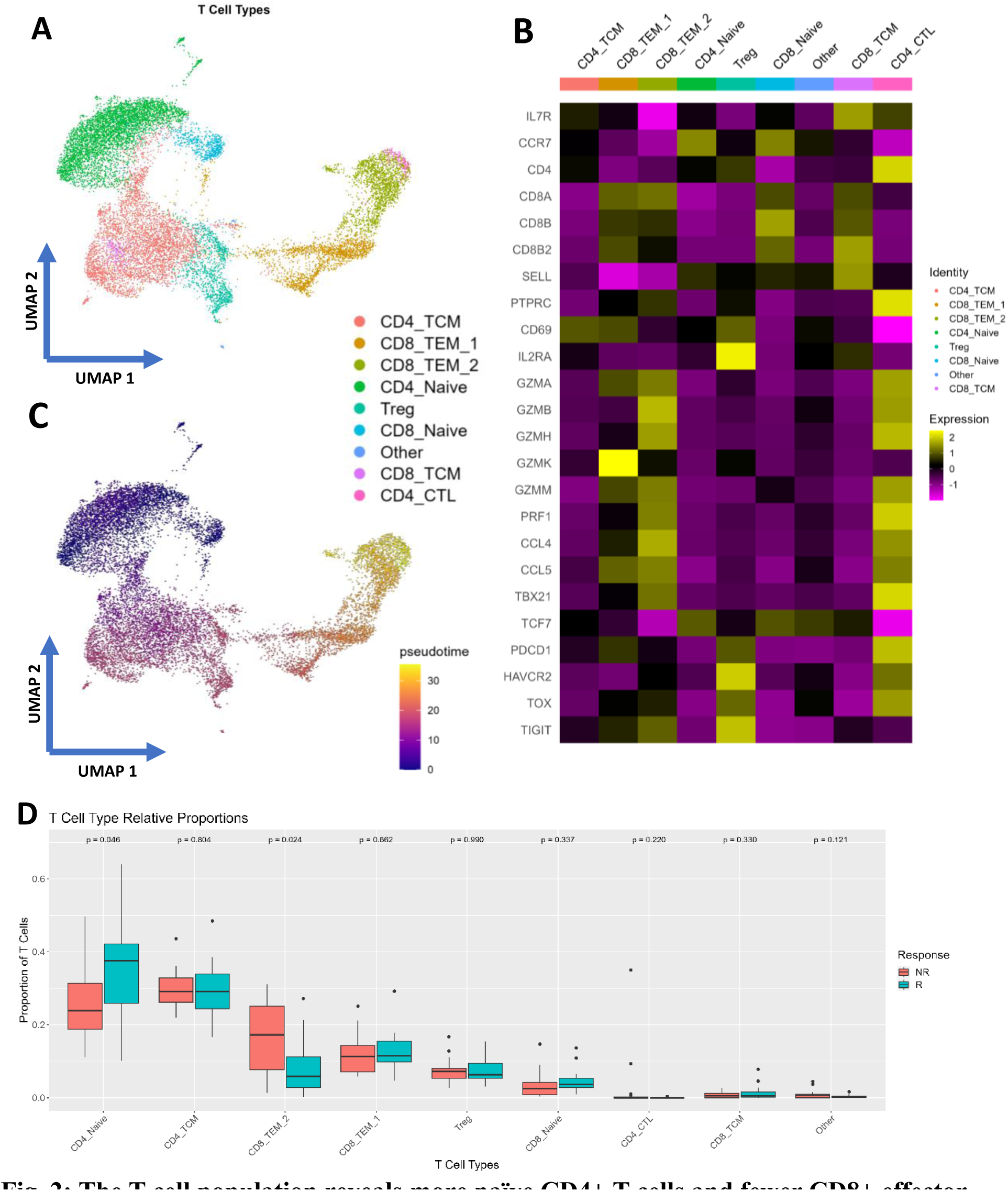
The T cell population reveals more naïve CD4+ T cells and fewer CD8+ effector memory T cells in responders pre-vaccination. **(A** and **B)** Uniform manifold approximation and projection (UMAP) integrated embedding of T cell subset, colored by (A) cell type and (B) Pseudotime **(C)** Marker gene signatures in T cell types (normalized averaged expression by cell type) **(D)** Paired box plot representing relative average proportion of T cell population in responders vs non-responders (p-values calculated using t-test)

Finally, analysis of the relative proportion of T cell sub-types in pre-vaccination PBMCs found that responders contained 34.7% more CD4+ naïve T cells, while non-responders showed 87.9% more CD8+ TEM group 2 cells (**Figure 2D**).

### Pre-vaccination, T cells of responders are at higher activation, transcription, and translation states than of non-responders

After identifying PBMC differences at the cellular level between responders and non-responders, we performed differential gene expression analysis in each individual T cell subtype and across the whole T cell population. This analysis revealed several consistently highly differentially expressed genes (DEGs) such as FOS and JUN, which are part of the AP-1 transcription factor (**Figure 3A**). We also found that the ribosomal protein RPS26, an important gene for T cell survival (*34*), was expressed higher in T cells of responders compared to non-responders. At the same time, expression of cytotoxic markers GNLY and NKG7 is consistently reduced. These genes that are differentially expressed at the single cell level are also seen differentially expressed at the pseudobulk level (**Figure 3B**).

**Fig. 3:**
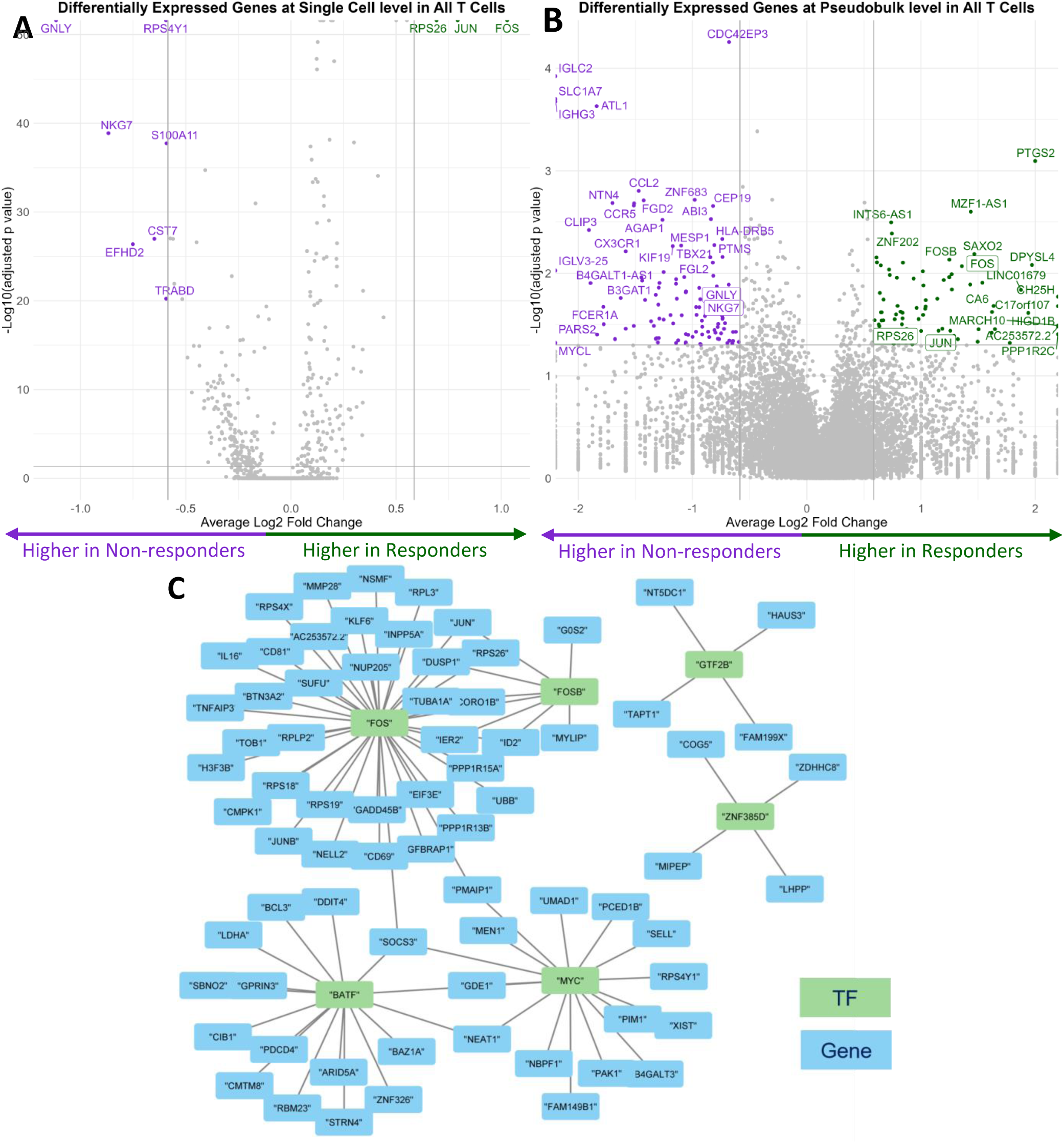
Differentially expressed gene analysis within the T cell populations show several core highly- and under-expressed gene signatures across all T cells and distinct expression profiles for different TEMs. **(A** and **B)** Differentially expressed genes at **(A)** single cell level (gene expressed in at least 25% of cells) and **(B)** pseudo-bulk level between response vs non-response across all T cells. **(C)** Scenic inferred transcriptional factors (Green) and their downstream targets (Blue) in T cells

We also inferred transcription factor expression in T cells from the gene expression data using SCENIC (*35*). Three inferred transcription factors, FOS, FOSB, and GTF2B, were positively correlated with immune response to the vaccine. The other three transcription factors, ZNF385D, BATF, and MYC, were negatively correlated with response (**Figure 3C**).

### Immune signaling pathways are enriched pre-vaccination in T cells of vaccine responders

Gene ontology (GO) (*36, 37*) and Reactome pathway (*38*) enrichment yielded higher translational and transcriptional activity in T cells of responders, shown in **Figure 4A and 4B**. Specifically, GO pathway enrichment found immune response pathways with higher immune and signaling receptor activity in T cells of responders compared to non-responders. Reactome also found, specifically in T cells, enriched calcium signaling and decreased antigen presentation. Results from Ingenuity pathway analysis (IPA) show many canonical pathways related to immune signaling expressed higher in T cells of responders (**Figure 4C**). These included enriched EIF2 signaling, mTOR signaling, and IL-10 signaling.

**Fig. 4:**
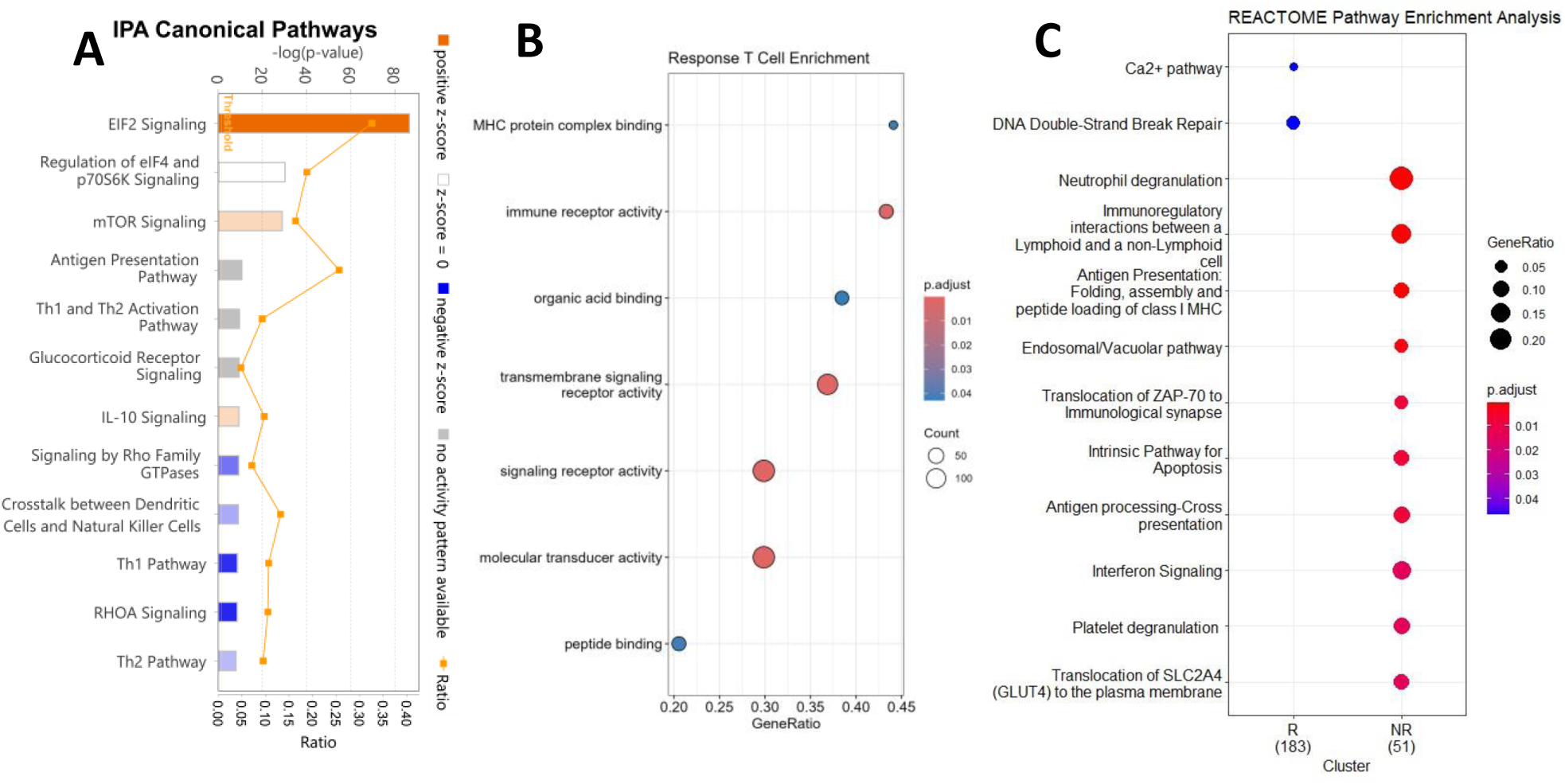
Pathway analysis of the T cell scRNA-seq data shows enriched immune signaling, immune cell cross talk, and decreased cell death in responders. **(A)** Go Pathway analysis **(B)** Reactome pathway analysis **(C)** IPA pathway analysis

### Network modeling reveals key gene signatures connected to vaccine response

The presence of pre-vaccination scRNA-seq features that differ significantly between vaccine responders and non-responders indicates that some of them may directly affect the response. To investigate this we applied directed (causal) graph network estimation using CausalMGM (*39, 40*). **Figure S6** represents the learned networks using all single cell data, while **Figure 5** represents the CD4+ naïve and CD8+ TEM Group 2 T cell networks built from the pseudobulk data. The edges in these graphs represent conditional dependencies (i.e., the two variables still share information for each other even after we account for any other variable or combinations of variables in the dataset); thus, the connections can be considered of representing direct effects.

**Fig. 5:**
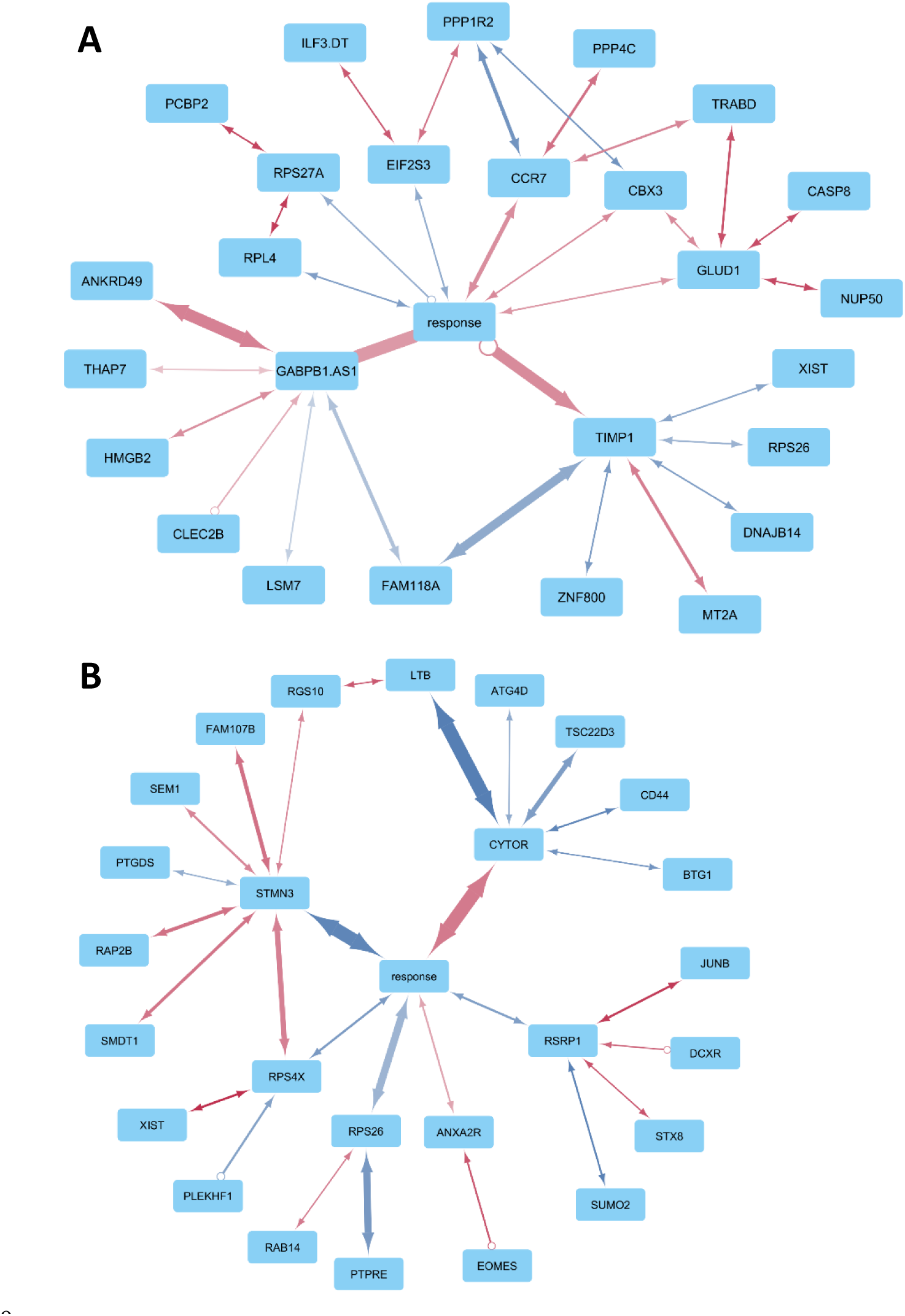
Network inference via rCausalMGM reveals possible gene interactions from pseudobulk T cell data. Networks for (**A**) CD4+ Naïve T Cells and (**B**) CD8+ TEM Group 2 inferred by rCausalMGM software using MGM undirected search and FCI-stable directed search. Edge color represents the correlation between the two features, with blue indicating positive, and red indicating negative correlation. Edge thickness represents the stability of the edge across cross validation, with thicker lines indicating that the edge appeared across an increased number of folds.

Figure S6A shows the first neighbors of response in the undirected MGM network. **Figure S6B** shows that there are 24 genes within second neighbors of response for CD8+ effector memory cells, a superset of the genes that isolates response from the rest of the graph (the Markov blanket). Across both single cell-based networks, we see several repeated genes of interest, such as FOS, JUN, and JUNB. In the TEM network, we also see the HLA complex represented in HLA-A and HLA-B, as well as many cytotoxic markers such as GZMM. We see CCL5 represented again through KLF2, a transcription factor that controls CCL5. We also see genes that regulate T cell differentiation and activity, such as TGFB1, and NFKBIA (which regulates NFKB).

In Figure 5, we see the ensemble Markov Blanket (*41*) of response extracted from the whole CausalMGM network generated from pseudobulk data with Leave-One-Out Cross-Validation (LOOCV). The stability of each edge across all folds can be seen by the width of the edge. For naïve CD4+ T cells, in the genes that occur most commonly in the Markov Blanket, we see GABPB1-AS1, ANKRD49, TIMP-1, and FAM118A. For CD8+ TEM group 2 cells, we see CYTOR, STMN3, and LTB.

### Response to the vaccine at week 12 can be predicted from pre-vaccination PBMC scRNA-seq data

We trained several different popular types of machine learning (ML) models and measured their performance at different parameter thresholds using LOOCV. For performance evaluation we used the area under the receiver operating characteristic curve (AUROC).

In Figure 6 we see the overall performance of 5 different ML models in predicting individual response from pseudobulk count values of the CD4+ naïve and CD8+ TEM 2 cell types. We found that for CD4+ naïve T cells that using the network model generated by MGM for feature selection combined with a general linear model has the best overall performance with an AUC of 0.727, while for CD8+ TEM group 2, the XGBoost model using all differentially expressed genes has the best overall performance at 0.938. We can also see that the CD8+ TEM 2 based models generally had higher average performance.

**Fig. 6:**
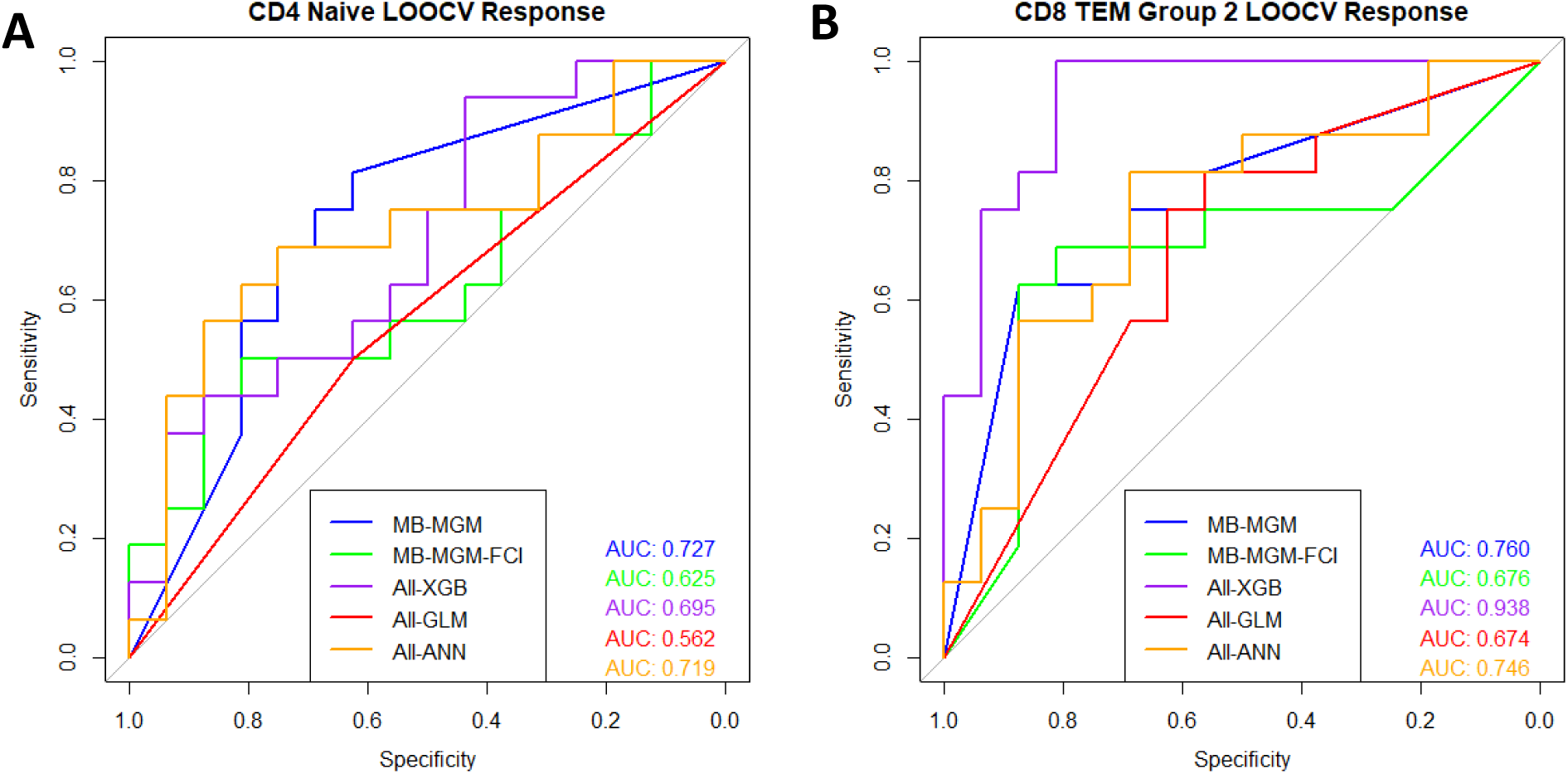
Predictive machine learning models can be built off pseudobulk data. Leave one out cross validation AUROC predictive performance of 5 ML models for (**A**) CD4+ naïve T cells and (**B**) CD8+ TEM Group 2. For MB-MGM and MB-MGM-FCI, the Markov blanket of the respective networks as seen in Figure 5, were used to select features used to train general linear models to do prediction. ALL-GLM is a general linear model trained on all gene features, while ALL-XGB and ALL-ANN are XGBoost and simple Neural Network models to serve as comparisons.

## DISCUSSION

In this paper, we demonstrated that scRNA-seq data have a clear pre-vaccination signal predictive of week-12 response. Specifically, responders had a significantly higher percentage of CD4+ naïve T cells while non-responders had a significantly higher percentage of terminally exhausted GZMB+ CD8+ TEM cells. A higher proportion of naïve T cells increases the likelihood and speed of priming of antigen-specific T cells, which was the goal of the preventative MUC1 vaccine. The effectiveness of the vaccine would be expected to be much lower in individuals with a higher number of TEM cells present with an exhausted phenotype.

There is also a higher CD16+ monocyte population in responders compared to non-responders. These “non-classical” monocytes are known to play a role in promoting an inflammatory response when a foreign antigen is detected in the peripheral blood (*42*). In preclinical studies in mouse models, non-classical monocytes have been shown to promote T cell activation (*43*) and to recruit regulatory T cells (*44*). The differences found in pre-vaccination PBMCs between responders and non-responders suggests that CD16+ monocytes play a role in priming the naïve CD4+ T cells to generate a response to the vaccine.

Focusing on gene expression, we see that JUN and FOS are more highly expressed across all T cell subtypes in responders than in non-responders. These genes were also identified as key genes in determining response through gene interaction network inference. The JUN and FOS family of genes are associated with the AP-1 transcription factor (TF), which was also inferred through SCENIC. The AP-1 TF plays an important role in T cell activation (*45*) and low AP-1 levels are associated with T cell exhaustion and anergy (*46–48*). RPS26 is also shown to be highly overexpressed in responders. Previous research shows that this ribosomal protein is a critical checkpoint for T cell survival and homeostasis in a p53-dependent manner (*34*). When RPS26 is knocked out, peripheral T cell homeostasis is impaired and T cell development is arrested in the thymus.

In comparison, T cells generally demonstrated lower levels of GNLY and NKG7 in responders, which has been associated with the GZMB+ TEM population both in our data and in other studies (*49*). Higher levels of GNLY, NKG7, and GZMB as well as the BATF TF are associated with terminally differentiated effector T cells, which agrees with differences we found in relative proportions of T cell subtypes. MYC TF, which is downregulated in responders, is more highly expressed in already activated T cells, but less expressed in quiescent naïve T cells (*50*). Like RPS26, CCL5 is responsible for regulating immune response and immune cell homeostasis. In mouse studies, deletion of CCL5 enhancers results in higher cytotoxic activity of T and NK cells and lower tumor metastasis, while higher levels are associated with increased metastasis (*51*).

Pathway analysis found enriched immune response pathways, with higher immune and signaling receptor activity in responders than in non-responders. The enriched adaptive immune response pathways indicates that responders are better prepared to respond, i.e. “ready-to-go”, while non-responders show lower levels of “readiness” of these pathways. Responders had enrichment in calcium signaling but low antigen presentation activity, indicating the presence of T cell populations ready to be primed, and no evidence of ongoing T cell differentiation and clonal expansion. Several additional pathways enriched in responders included EIF2 signaling and immune cell crosstalk. EIF2 signaling, which is downstream of T cell activation, inactivates Glycogen Synthase Kinase-3 (*52*). This can have many downstream effects, including blocking PD-1 expression, which in turn enhances T cell cytotoxic response (*53*). Crosstalk signaling included mTOR signaling between T cells and DCs, which drives naïve T cell differentiation into Th cells (*54*), and IL-10 signaling, which can have an immunosuppressive role (*55*).

We performed pseudobulk analysis for each individual T cell subpopulation and built predictive models on the cell type specific pseudobulk datasets. In both the CD4+ naïve and CD8+ TEM, we were able to build models with high predictive performance, with the best performing models for each population having AUC=0.727 and AUC=0.938, respectively. This indicates that there are clear pre-vaccination differences within the different T cell subpopulations that can be used to predict at a relatively high success rate who will and who will not respond to the MUC1 vaccine. This may also lead us to ways in which we might convert predicted non-responders into responders.

The advantage of network based predictive models is that we can see the genes that are important for the response in the LOOCV networks. For the CD4+ naïve T cell population, 4 main genes are present in most cross-validation folds. GABPB1-AS1 is negatively associated with response and is a long non-coding RNA (lncRNA) that regulates GABPB1. Upregulation of GABPB1-AS1 can eventually suppress the cellular antioxidant capacity and has been found to serve as a potential biomarker in prostate cancer (*56–58*). ANKRD49 is positively associated with response, and is responsible for upregulating cell proliferation (*59, 60*). TIMP-1 is negatively associated with response and is an important regulator of inflammatory processes (*61–63*). Elevated levels of TIMP-1 generally correlate with the progression and poor prognosis of various diseases. Finally, FAM118A is negatively associated with response and is predicted to be integral component of cell membranes and may play a role in T cell function in skin disease (*64–66*).

For the CD8+ TEM group 2 networks, there are 3 genes that are in the majority of folds. The CYTOR lncRNA is an oncogene that is negatively associated with response, and is related to immune pathways, aberrant glycolysis, and mitochondrial respiration. CYTOR plays a role in immune cell infiltration and is positively correlated with PD-1 expression; increased expression of CYTOR correlates with worse overall survival in cancer (*67–69*). STMN3 is negatively correlated with response and plays an important in anergic TTT cells as well as in signal transduction, causing cellular proliferation defects and cell cycle arrest when overexpressed (*70*). Finally, LTB is negatively associated with response and has a role in protective TTT cell response, TTT cell differentiation, and regulating T cell homeostasis (*71–73*).

In summary, we were able to determine specific cell types and gene signatures that are different pre-vaccination between immune responders and non-responders to the vaccine. Using this information, we were able to develop accurate machine-learning models that predict vaccine response from pre-vaccination PBMC scRNA-seq data. We were also able to extract information from these models to support selection of candidate biomarkers of response to the MUC1 vaccine and likely other cancer vaccines and immunotherapies, for future development and validation.

## MATERIALS AND METHODS

All descriptions of Materials and Methods should be included in the main paper. The Materials and Methods should be broken up into sections, each with a short subheading. Please include a study design paragraph at the start of the Materials and Methods and a statistical paragraph at the end of the Materials and Methods in the main text. If the Materials and Methods make the paper exceed the length limitations, less important sections of the Materials and Methods can be moved to the Supplementary Materials.

### Patient Data and Selection

This study selected PBMCs from 16 responders and 16 non-responders to the MUC1 vaccine, with response determined by ratio of anti-MUC1 IgG titer at week 0 pre-vaccination and week 12 post-vaccination, from two Phase II trials, NCT007773097 (*21*) and NCT02134925 (*22*),. The participants were individuals with a history of advanced colonic adenoma diagnosis who were at elevated risk for developing new polyps and an increased risk for colon cancer. The first trial tested immunogenicity and safety of the vaccine and the second trial also tested the vaccine efficacy, the ability to prevent new adenoma formation. PBMC and plasma were frozen and banked for future analysis. Associated demographic and clinical data included age, gender, ethnicity, and BMI.

### Approval Information

The NCT007773097 study was approved by the University of Pittsburgh Institutional Review Board and monitored by Data Safety Monitoring Board of the Clinical Translational Science Institute of the University of Pittsburgh (Pittsburgh, PA). The vaccine was approved by the U.S. Food and Drug Administration (FDA) as an Investigational New Drug (IND). The NCT02134925 study was approved by the Institutional Review Board at each of the six involved clinical centers: the Mayo Clinic, Rochester MN; the University of Pittsburgh Medical Center, Pittsburgh PA; the University of Puerto Rico, San Juan PR; the Veterans Administration, Kansas City, KS; Thomas Jefferson University, Philadelphia PA; and Massachusetts General Hospital, Boston MA. The National Cancer Institute Division of Cancer Prevention administratively coordinated the trial through The Cancer Prevention Network at the Mayo Clinic. Written informed consent was obtained from all participants across both studies.

### Peripheral Blood Mononuclear Cells (PBMCs)

All blood samples from both trials were processed and stored using the same protocol by the same operator. Each sample was processed within 24 hours of being drawn. The heparinized blood was layered on the lymphocyte separation medium (MPBio) and centrifuged at 800 g for 10 min at the lowest acceleration and deceleration speed. PBMCs were then collected from the interphase between the plasma and separation medium, washed twice, and resuspended in 80% human serum and 20% DMSO. The PBMCs were then sub-aliquoted to ∼10^7 cell vials, cryopreserved, and stored in liquid nitrogen until use.

### Single Cell RNA Sequencing

Single cell RNA-sequencing (scRNA-seq) was performed on the banked pre-vaccination PBMCs in four batches, each with 8 individuals. The target sequencing depth was 2000 cells per patient, with 50000 reads per cell. Two individuals processed the initial two batches in tandem following the same protocol for 3’v2 sequencing, and two individuals processed the second two batches in tandem following the same protocol for 5’v2 sequencing. All sequencing was performed using 10x Genomics technology by the Single Cell Core at the University of Pittsburgh.

## 3’v2 Sequencing

The first two batches were done in one sequencing experiment with 3’ v2 sequencing technology. Banked PBMC samples were thawed into warm complete Roswell Park Memorial Institute (cRPMI) medium. Then, the samples were pelleted, washed, and resuspended in cRPMI to remove any remaining DMSO. The viability of each sample was measured before the samples were pelleted, washed, and resuspended in staining buffer (2% FBS, 0.01% Tween, in PBS).

Then Fc Blocking Reagent was added to the samples and incubated for 10 minutes at 4 °C. Next, the hashing antibodies (*74*) were added to the samples and incubated for 30 minutes at 4 °C to bind. The samples were then washed, resuspended in 0.04% BSA in PBS, and filtered through 40 um strainers. The viability of the cells was measured again before the samples were combined into two batches of 8 patients. The mixed samples were then sequenced using the 3’ v2 protocol using the Illumina NextSeq-500 through the University of Pittsburgh Single Cell Core as described in previous papers (*75, 76*).

### 5’v2 Sequencing

The second sequencing experiment was done with 5’ v2 sequencing technology. Banked PBMC samples were thawed and mixed into warm cRPMI. The samples were pelleted, washed, and resuspended in cRPMI to remove any remaining DMSO. The viability of each sample was measured before the samples were pelleted, washed, and resuspended in staining buffer (2% FBS, 0.01% Tween, in PBS). The hashing antibodies were diluted with PBS, before being combined with the corresponding samples and incubated for 30 minutes at 4 °C to bind. The samples were then washed 3 times and resuspended in staining buffer. The viability of the cells was measured again before the samples were combined into two batches that were suspended in 0.04% BSA in PBS. The hash tagged batches were then sequenced using the Illumina 5’ v2 using the Illumina NextSeq-500 through the University of Pittsburgh Single Cell Core as described in previous papers (*77*).

### Mapping and Alignment

Raw FASTQ read files were run through Cell Ranger (10x Genomics Cell Ranger 5.0.1) (*78*) in order to align reads to HG38 reference genome to generate count .h5 files, which include gene and hashtag information, which were then loaded into R as count matrices using Seurat v4 (*33*).

### Bioinformatic and ML Analysis

All downstream analysis were done in R <= v4.3.1, Python <= 3.10.12, and Julia <= 1.6, with the exact environment versions dependent on library and package requirements for individual analyses pipelines.

### Clustering

To find the cell types within the sequenced cell population, the integrated data was clustered. The clustering was done using a shared nearest neighbor (SNN) modularity optimization based clustering algorithm (*79*), available through Seurat through its FindNeighbors and FindClusters functions. The SNN was calculated using the 100 nearest neighbors (other parameters were default). In order to determine the number clusters to find, a hyperparameter search was done to determine cluster stability at different resolution values. The greatest resolution with no unstable clusters (no cell movement between clusters) was selected.

The clusters were then annotated manually and automatically. For manual annotation, the top expressed and top differentially expressed genes (DEG) in each cluster were compared to known marker genes for PBMC cell types. For automated annotation, the dataset was mapped to an existing multi-modal PBMC dataset (*33*) using Seurat’s built in mapping pipeline (default parameters).

### Pseudotime Trajectory

In order to estimate the pseudotime and the cell trajectories, two main algorithms were used: (1) Monocle3 (*80–82*) and (2) scVelo (*83*). Both were run following the standard pipelines with default parameters. For Monocle3, we used Leiden (*84*), clustering (*85*), and partitioning (*85*).

### Comparative Analysis

To determine differences between responders and non-responders, differential gene expression (DGE), gene set enrichment, pathway analysis were run. Differential gene expression was done using the built in Seurat methods to find markers for specific conditions and clusters. The average expression of each gene for responders and non-responders was calculated for each cell type using Seurat’s AverageExpression function. This was used to generate a p-value ranked matrix of DEG using Wilcoxon rank sum and Bonferroni correction. Pathway analysis was run using ssGSEA, Reactome, and Ingenuity Pathway Analysis which allows canonical, gene set, and gene ontology pathway enrichment.

### Mechanisms and Networks

To estimate regulatory information from the single cell data, the regulon and transcription factors were inferred using SCENIC (*35*). This allowed us to estimate transcription factor and gene pairs, as well as build connectivity maps of the regulon. We also used CausalMGM to generate gene interaction networks.

CausalMGM is an algorithm developed to learn undirected and directed graphical models. A probabilistic graphical model is a graph/network where each node represents a feature, and each edge between two nodes represents a relationship between the two features given all the other features in the dataset. In a simple undirected graphical model, each edge would solely represent a conditional dependence. In a directed graphical model, the direction of the edge would indicate the direction of that relationship. A directed model relies on additional rules based on causal graph theory in order to orient the direction of each edge. The algorithm begins by inferring an undirected graphical model using the mixed graphical model algorithm (MGM), and then using a constraint-based algorithm to further refine the edges and estimate edge directions. The multi-step process has been implemented in R and is available on GitHub at https://github.com/tyler-lovelace1/rCausalMGM.

### Prediction Models and Evaluation

Two main strategies were tested in the prediction models. First, the scRNA-seq data of each target cell population was used directly to train the models. Second, pseudobulk values were calculated for each individual for each target cell population was used to train the models. Pseudobulk count values are a cumulative sum of all the single cell counts for a specific population.

Two main sets of features were used to train the models. First, all the DEG between responders and non-responders for the specific cell population of interest were used as the baseline feature set. Second, networks were generated by the CausalMGM algorithm and then the genes that are conditionally dependent with the response variable (the Markov blanket of response) were selected as a reduced feature set.

Several popular machine learning algorithms were tested to develop prediction models. These were the generalized linear models (GLM), XGBoost from XGBoost R package, and simple neural networks from the NeuralNet R package.

To evaluate the performance of the predictive model on the single cell count dataset, two main types of cross validation (CV) were performed. These were 10-fold stratified CV, and leave two patients out CV (one responder, one non-responder, randomized pairs). For the pseudobulk count dataset, leave one out cross validation (LOOCV) was performed (which is the same leave one patient out in this specific scenario). The probability thresholds from each test split were recorded, and a Receiver Operating Characteristic (ROC) curve was generated for each algorithm and dataset. The area under the curve (AUC) was calculated to determine the performance of the models. All related statistical and machine learning related analysis was performed in R.

## Acknowledgments

We thank Dr. R. Schoen, PI of both trials, and the NCI DCP Cancer Prevention Network (CPN) for support of the multi-center trial. We appreciate the tremendous efforts of the team members that conducted the two clinical trials and provided samples for this study, particularly Lisa Boardman, Marcia Cruz-Correa, Ajay Bansal, David Kastenberg, Chin Hur, Lynda Dzubinski, Sharon Kaufman, Luz M Rodriguez, Ellen Richmond, Asad Umar, Eva Szabo, Andres Salazar, Ryan McMurray, Carrie Strand, Nathan Foster, David Zahrieh, and Paul Limburg. We are also indebted to all the trial participants for their commitment to our cancer prevention mission. We thank Dr John McKolanis for processing the PBMCs, Matthew T. Dracz, and Najla Saadallah for technical support in preparing cells for sequencing experiments, and Tracy Tabib and the Single Cell Sequencing Core of the University of Pittsburgh School of Medicine for performing the sequencing.

## Funding

National Institutes of Health grant R01HL159805 (PVB) National Institutes of Health grant T32CA082084 (DYY) National Cancer Institute grant R35CA210039 (OJF)

## Authors’ Disclosures

OJF is member of the Scientific Advisory Boards of PDS Biotech, GeoVax and Invectys

## Data and materials availability

Code and data used to generate the figures can be found on GitHub at: https://github.com/benoslab/MUC1-Cancer-Vaccine-Single-Cell-RNAseq

## Supplementary Materials

### Materials and Methods

#### Quality Control and Normalization

Using Seurat pipeline, the two generated datasets for the two sequencing experiments were first filtered for low quality cells. Quality was determined by fraction of mitochondrial counts per cell (<5%) and number of features per cell (>200 features). Each run was demultiplexed into the original samples based on HTO enrichment using Seurat’s HTODemux function with default parameters. Negative and doublets were removed from the dataset. After quality control, each sample was normalized using sctransform (*86*)

#### Dimensionality Reduction

In order to visualize the data in 2D, dimensionality reduction was run on the batch-corrected dataset. Three types of dimensionality reduction were done for visualization: PCA, TSNE, and UMAP. Previous research has demonstrated that uniform manifold approximation and projection (UMAP) demonstrates the highest stability and best separation between the original cell populations (*87*). In order to generate the UMAP reduction, first PCA was run to generate the top 100 components. Then UMAP dimensions were calculated using the initial PCA reduction with all components.

#### Integration and Batch Correction

After quality control and normalization, the samples were integrated for batch correction. Two different types of integration were used: (1) Seurat’s built in integration using RPCA and (2) a variational autoencoder (VAE) based method SCVI (1.0.2) (*88*).

For RPCA based integration, the top 3000 highly variable genes were selected as to build the anchor set. Then the samples were integrated using these features via Seurat’s IntegrateData function run with default parameters (with SCT normalization).

For SCVI based integration, the top 5000 highly variable genes were used to train the VAE models, with batch number, patient number, and mitochondrial count fraction as covariate keys. The hyperparameters for the final model were selected using the built in raytune based hyperparameter search. After the training the initial VAE model, the annotated cell types can be used to build an extended model with scANVI (*89*) that has more accurate integration.

The integration methods were compared using scib-metrics (*90*) to determine the how well each method was able to perform batch correction while conserving biological information.

### Supplementary Figures and Results

#### Quality Control

scRNA-seq data were collected from samples directly before the first vaccination. The initial Cell Ranger alignment and aggregation identified a total of 34,234 genes over 45,729 cells with a total of 2,275,406,768 reads, with each individual batch seen in Figure S1. After initial antibody tagging to determine cell patient origin and filtering out low quality cells (mitochondrial read percentage < 5%, number of genes per cell > 200, doublets, and negatives), the final raw count matrix contained 22,888 genes across 38,224 cells from all 32 individuals.

#### Dimensionality Reduction

Several different types of dimensionality reduction algorithms were tested on the dataset. We can see the PCA, t-SNE, and UMAP reductions in Figure S2. We selected UMAP for later visualizations. The algorithms for these reductions can be found with the respective Seurat library (in R) and scverse ecosystem (in python).

#### Integration and Batch Correction

We ran two types of integration, Seurat and VAE-based (SCVI and scANVI). The integrated (batch corrected) values can be used to generate new UMAP dimensions. These can be seen in Figure S3A, B, C and show that the integration methods are able to correct for batch relatively well. In Figure S3D, we show the relative distribution of the different cell phases in the integrated dataset as a quality check to show that we aren’t generating cell cycle specific clusters.

#### Deconvolution

The initial exploratory study performed bulk RNA-sequencing. In order to utilize this data, bulk deconvolution was run using the single cell data as a reference. Instaprism (*91*) performs fast Bayesian based cell type deconvolution, and was used with default parameters to generate estimated cell type from the bulk RNA-seq data.

---

**Fig. S1:**
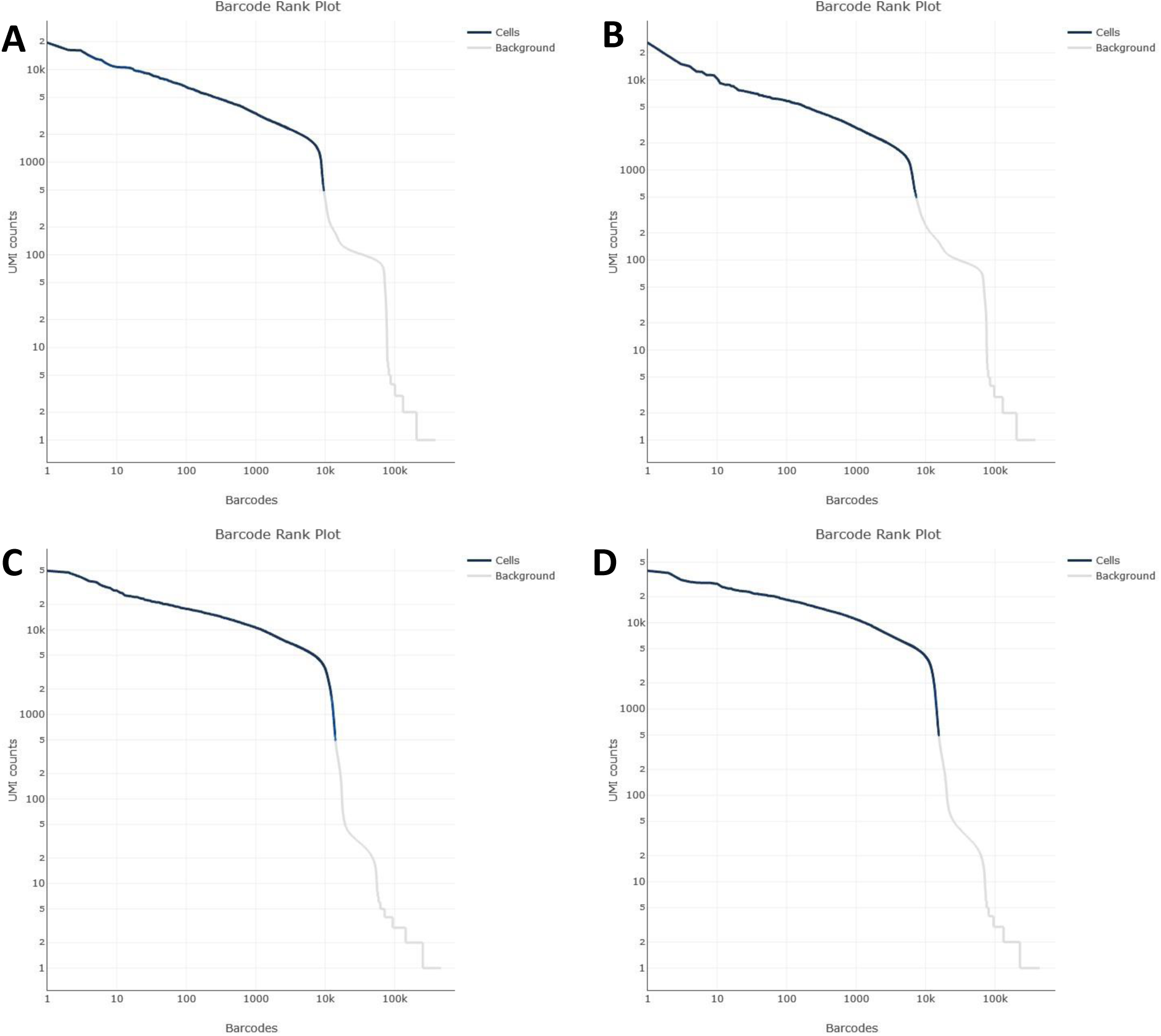
Observed transcript (UMI counts) per cell measured (barcodes) across each batch. **(A)** Experiment 1 Batch 1 **(B)** Experiment 1 Batch 2 **(C)** Experiment 2 Batch 1 **(D)** Experiment 2 Batch 2

**Fig. S2:**
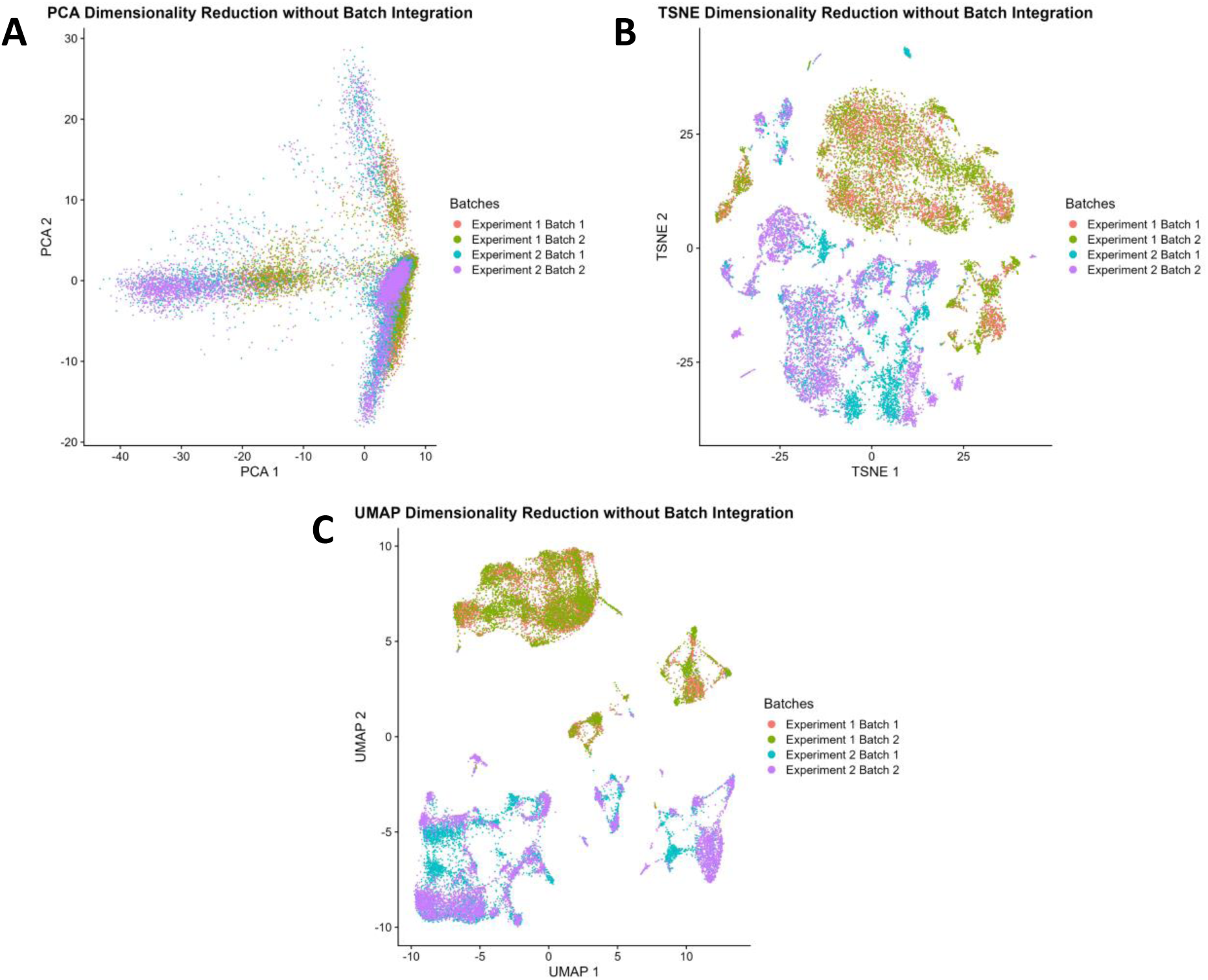
Dimensionality reduction without any batch correction. **(A)** PCA reduction **(B)** TSNE reduction **(C)** UMAP reduction

**Fig. S3:**
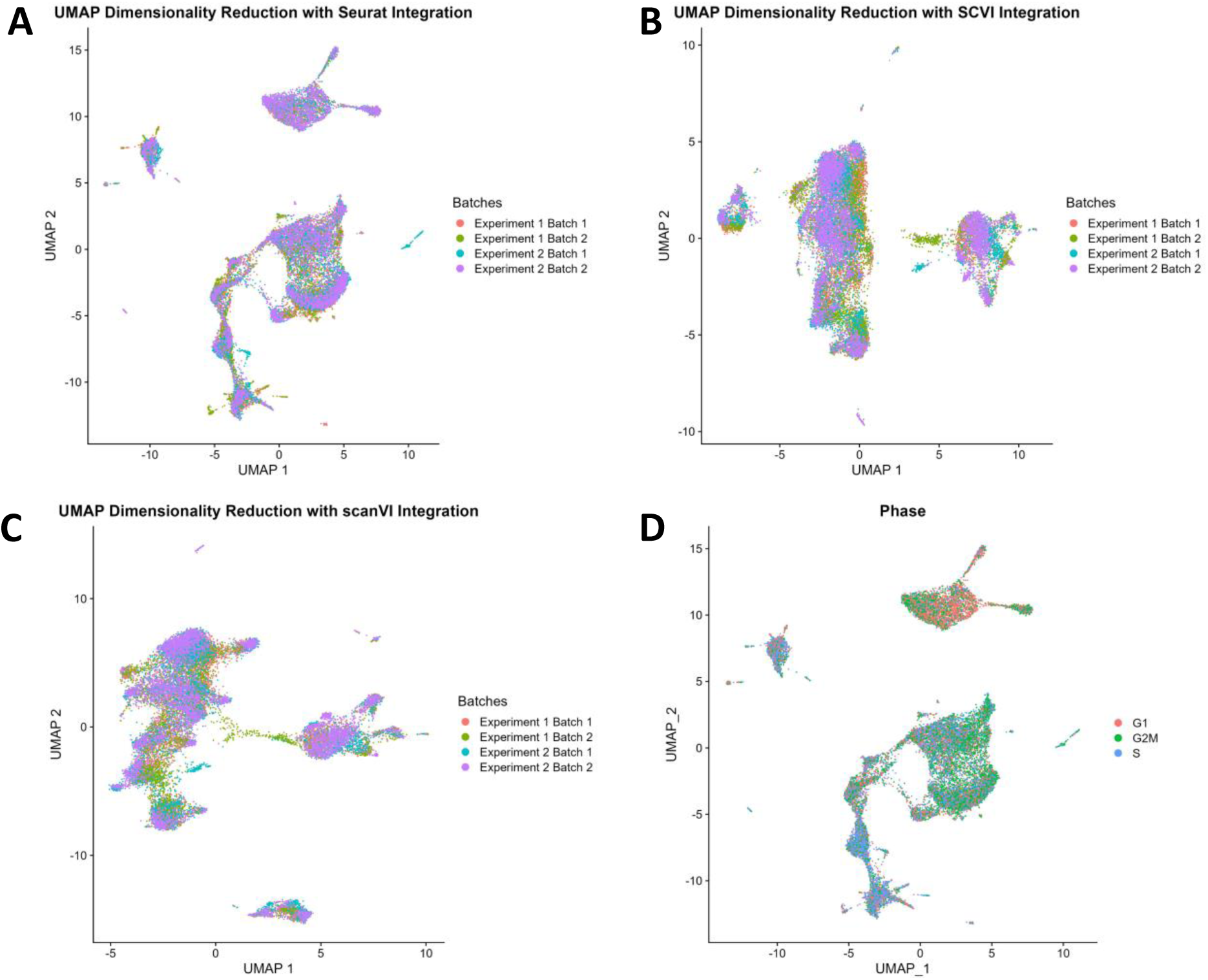
UMAP reduction with batch correction. **(A)** Seurat Integration **(B)** SCVI Latent **(C)** SCANVI Latent **(D)** Checking to make sure cell cycle is spread throughout UMAP reduction

**Fig. S4:**
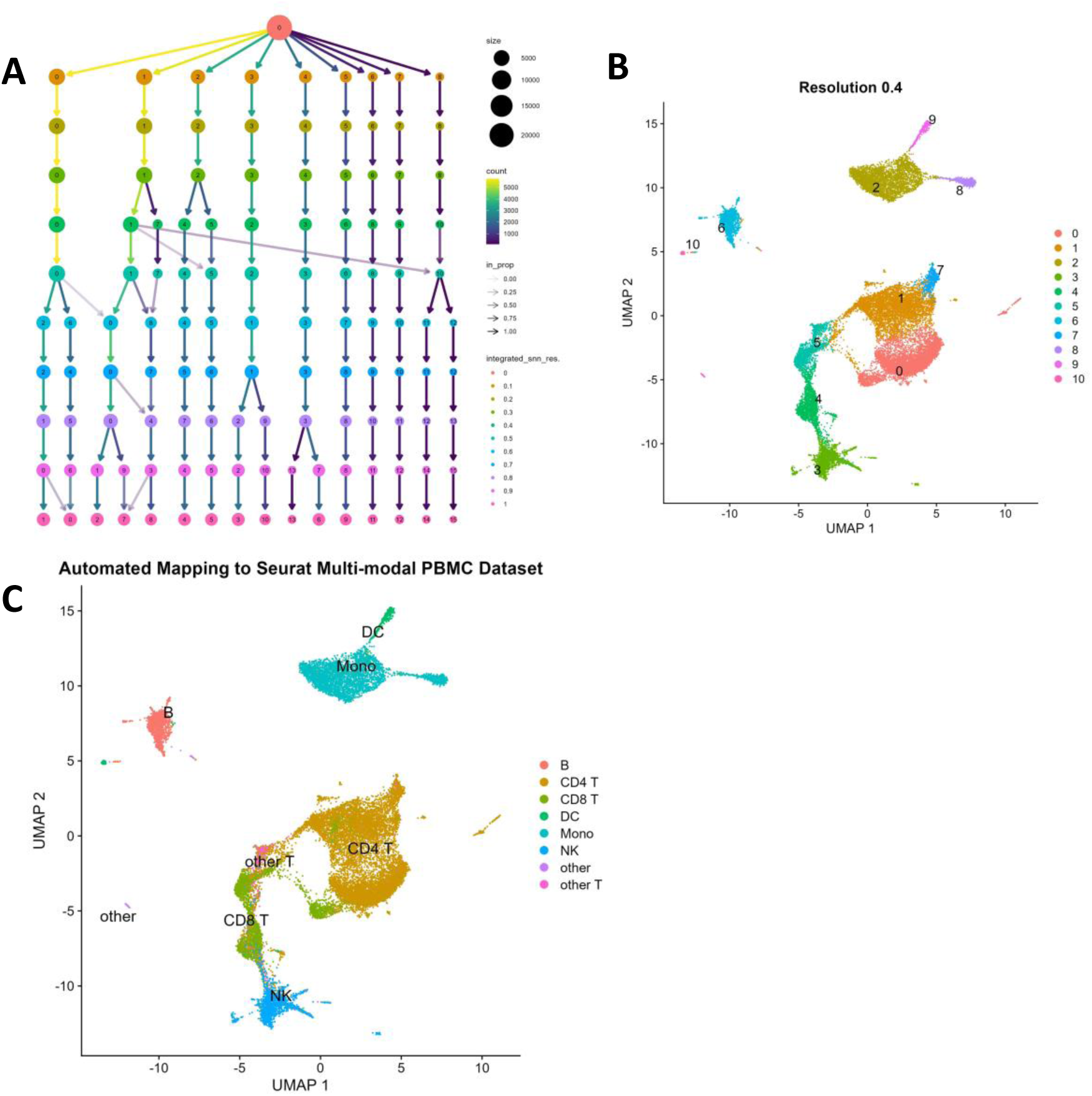
Determining the most stable clustering resolution by using the highest resolution without cluster membership changing. **(A)** Clustering membership over different resolutions **(B)** Clustering at selected resolution **(C)** Automated mapping of PBMC dataset onto existing Seurat reference

**Fig. S5:**
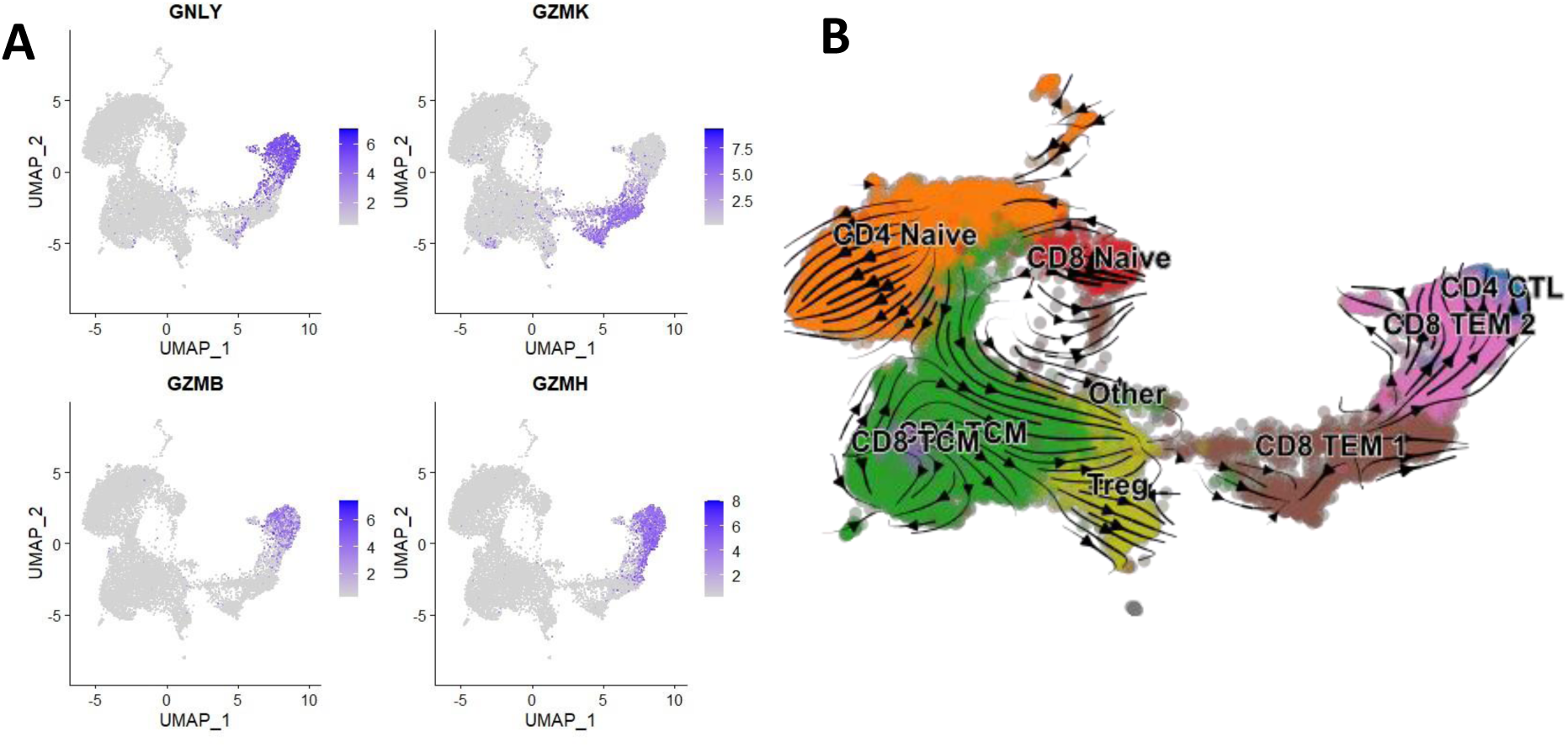
Additional figures for T cells. (A) Granzyme and cytotoxic gene expression in T cells (B) Estimated RNA velocity in T cells

**Fig. S6:**
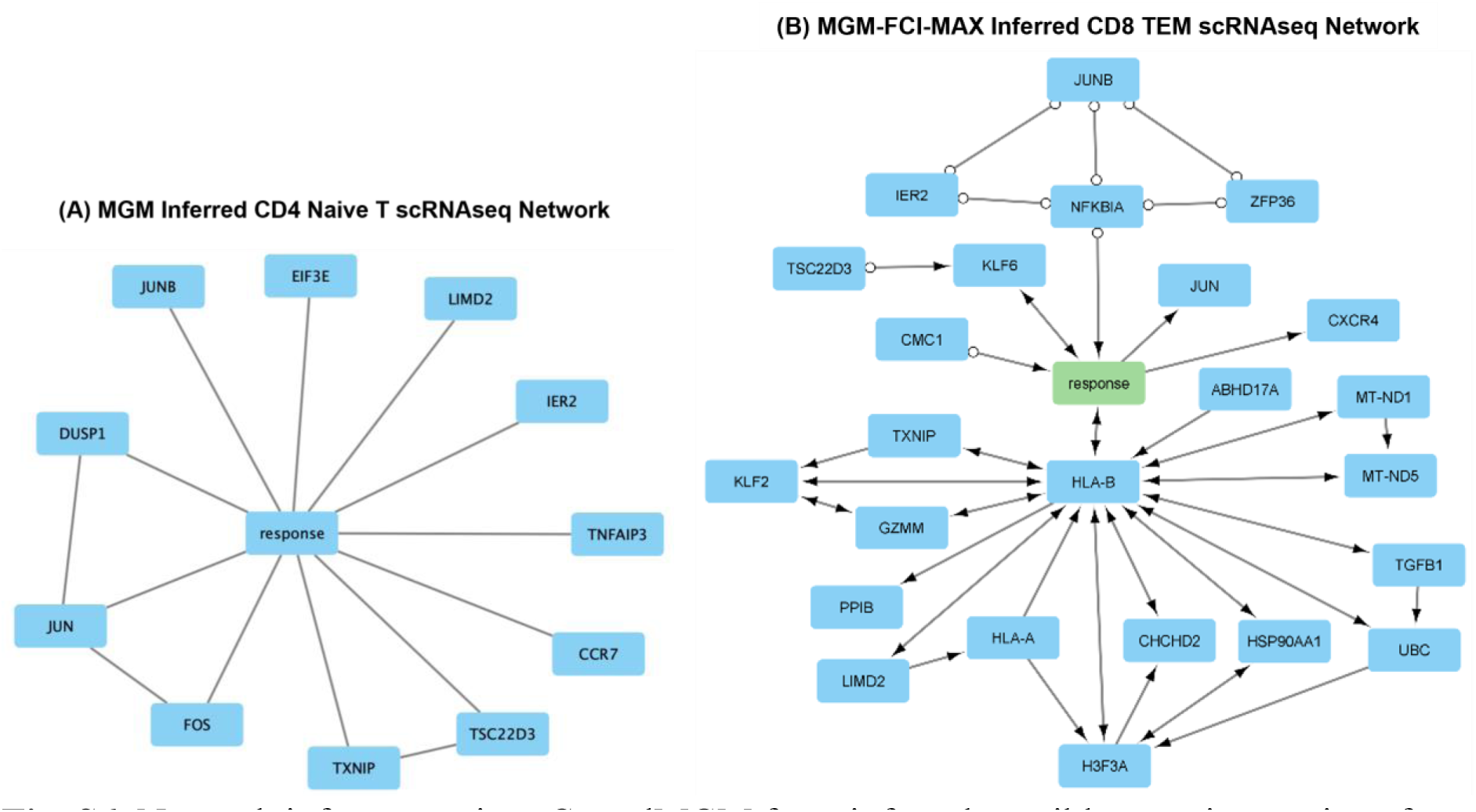
Network inference using rCausalMGM from inferred possible gene interactions from single cell data pre-vaccination. **(A)** Predicted undirected gene-response network for Naïve CD4+ T cells using MGM **(B)** Predicted directed gene-response network CD8+ TEM using MGM-FCI-MAX

## Notes

### Summary of Updates

Minor revisions to main body text for clarity, updated funding, added short summary, updated author correspondence information

## References

1. R. American Association for Cancer, AACR Cancer Progress Report 2023.

2. P. Sharma, J. P. Allison, Dissecting the mechanisms of immune checkpoint therapy. Nature Reviews Immunology 20, 75–76 (2020).

3. S. K. Nakhoda, A. J. Olszanski, Addressing recent failures in immuno-oncology trials to guide novel immunotherapeutic treatment strategies. Pharmaceutical medicine 34, 83--91 (2020).

4. J. Bach, K. Wojas-Krawczyk, Nico, Failure of immunotherapy—the molecular and immunological origin of immunotherapy resistance in lung cancer. International Journal of Molecular Sciences 22, 9030 (2021).

5. M. Karasarides et al., Hallmarks of resistance to immune-checkpoint inhibitors. Cancer immunology research 10, 372–383 (2022).

6. E. Balta, G. H. Wabnitz, Y. Samstag, Hijacked Immune Cells in the Tumor Microenvironment: Molecular Mechanisms of Immunosuppression and Cues to Improve T Cell-Based Immunotherapy of Solid Tumors. International Journal of Molecular Sciences 22, 5736 (2021).

7. Z. Ren, X. Zhang, Y.-X. Fu, Facts and hopes on chimeric cytokine agents for cancer immunotherapy. Clinical Cancer Research, (2024).

8. U. Uslu, C. H. June, T-cell Therapies Targeting Multiple Cancer Antigens: The Power of Many. Cancer Immunology Research, OF1-OF1 (2023).

9. M. D. Blunt, S. I. Khakoo, Harnessing natural killer cell effector function against cancer. Immunotherapy Advances 4, ltad031 (2024).

10. K. Tsuchikama, Y. Anami, S. Y. Y. Ha, C. M. Yamazaki, Exploring the next generation of antibody–drug conjugates. Nature Reviews Clinical Oncology, 1–21 (2024).

11. Z. Hu, P. A. Ott, C. J. Wu, Towards personalized, tumour-specific, therapeutic vaccines for cancer. Nature Reviews Immunology 18, 168--182 (2018).

12. O. J. Finn, The dawn of vaccines for cancer prevention. Nature Reviews Immunology 18, 183--194 (2018).

13. S. V. S. Deo, J. Sharma, S. Kumar, GLOBOCAN 2020 Report on Global Cancer Burden: Challenges and Opportunities for Surgical Oncologists. Annals of Surgical Oncology 29, 6497–6500 (2022).

14. O. J. Finn, Human tumor antigens yesterday, today, and tomorrow. Cancer immunology research 5, 347–354 (2017).

15. M. A. Cheever et al., The prioritization of cancer antigens: a national cancer institute pilot project for the acceleration of translational research. Clinical cancer research 15, 5323–5337 (2009).

16. O. J. Finn, Cancer vaccines: between the idea and the reality. Nature Reviews Immunology 3, 630–641 (2003).

17. M. E. Janes, A. P. Gottlieb, K. S. Park, Z. Zhao, S. Mitragotri, Cancer vaccines in the clinic. Bioengineering & Translational Medicine, (2023).

18. O. J. Finn, Premalignant lesions as targets for cancer vaccines. The Journal of experimental medicine 198, 1623–1626 (2003).

19. N. Çuburu, O. J. Finn, S. H. Van Der Burg, Cancer Prevention: Targeting Premalignant Epithelial Neoplasms in the Era of Cancer Immunotherapy and Vaccines. Frontiers in immunology 13, 924099 (2022).

20. T. Kimura, O. J. Finn, MUC1 immunotherapy is here to stay. Expert Opinion on Biological Therapy 13, 35–49 (2013).

21. T. Kimura et al., MUC1 vaccine for individuals with advanced adenoma of the colon: a cancer immunoprevention feasibility study. Cancer prevention research 6, 18--26 (2013).

22. R. E. Schoen et al., Randomized, Double-Blind, Placebo-Controlled Trial of MUC1 Peptide Vaccine for Prevention of Recurrent Colorectal Adenoma. Clinical Cancer Research 29, 1678--1688 (2023).

23. J. S. Goydos, E. Elder, T. L. Whiteside, O. J. Finn, M. T. Lotze, A phase I trial of a synthetic mucin peptide vaccine: induction of specific immune reactivity in patients with adenocarcinoma. Journal of Surgical Research 63, 298–304 (1996).

24. R. K. Ramanathan et al., Phase I study of a MUC1 vaccine composed of different doses of MUC1 peptide with SB-AS2 adjuvant in resected and locally advanced pancreatic cancer. Cancer Immunology, Immunotherapy 54, 254–264 (2005).

25. A. J. Lepisto et al., A phase I/II study of a MUC1 peptide pulsed autologous dendritic cell vaccine as adjuvant therapy in patients with resected pancreatic and biliary tumors. Cancer therapy 6, 955 (2008).

26. E. Scheid et al., Tn-MUC1 DC vaccination of rhesus macaques and a phase I/II trial in patients with nonmetastatic castrate-resistant prostate cancer. Cancer immunology research 4, 881–892 (2016).

27. T. J. Tan et al., A phase I study of an adenoviral vector delivering a MUC1/CD40-ligand fusion protein in patients with advanced adenocarcinoma. Nature Communications 13, (2022).

28. S. Ota, M. Miyashita, Y. Yamagishi, M. Ogasawara, Baseline immunity predicts prognosis of pancreatic cancer patients treated with WT1 and/or MUC1 peptide-loaded dendritic cell vaccination and a standard chemotherapy. Human Vaccines & Immunotherapeutics 17, 5563–5572 (2021).

29. M. Bilusic et al., Phase I study of a multitargeted recombinant Ad5 PSA/MUC-1/brachyury-based immunotherapy vaccine in patients with metastatic castration-resistant prostate cancer (mCRPC). Journal for immunotherapy of cancer 9, (2021).

30. C. C. Schimanski et al., Adjuvant MUC vaccination with tecemotide after resection of colorectal liver metastases: a randomized, double-blind, placebo-controlled, multicenter AIO phase II trial (LICC). OncoImmunology 9, 1806680 (2020).

31. M. Antonilli et al., Triple peptide vaccination as consolidation treatment in women affected by ovarian and breast cancer: Clinical and immunological data of a phase I/II clinical trial. International Journal of Oncology 48, 1369–1378 (2016).

32. C. M. Cameron et al., Pre-vaccination transcriptomic profiles of immune responders to the MUC1 peptide vaccine for colon cancer prevention. *medRxiv*, 2024.2005.2009.24305336 (2024).

33. Y. Hao et al., Integrated analysis of multimodal single-cell data. Cell 184, 3573--3587 (2021).

34. C. Chen et al., Ribosomal protein S26 serves as a checkpoint of T-cell survival and homeostasis in a p53-dependent manner. Cellular \& Molecular Immunology 18, 1844--1846 (2021).

35. S. Aibar et al., SCENIC: single-cell regulatory network inference and clustering. Nature methods 14, 1083--1086 (2017).

36. M. Ashburner et al., Gene Ontology: tool for the unification of biology. Nature Genetics 25, 25–29 (2000).

37. C. The Gene Ontology et al., The Gene Ontology knowledgebase in 2023. Genetics 224, iyad031 (2023).

38. A. Fabregat et al., The Reactome Pathway Knowledgebase. Nucleic Acids Research 46, D649–D655 (2018).

39. A. J. Sedgewick, J. Ramsey, P. Spirtes, C. Glymour, P. V. Benos, Mixed Graphical Models for Causal Analysis of Multi-modal Variables. *ArXiv* abs/1704.02621, (2017).

40. X. Ge, V. K. Raghu, P. K. Chrysanthis, P. V. Benos, CausalMGM: an interactive web-based causal discovery tool. Nucleic acids research 48, W597–W602 (2020).

41. C. F. Aliferis, I. Tsamardinos, A. Statnikov. (American Medical Informatics Association), vol. 2003, pp. 21.

42. P. B. Narasimhan, P. Marcovecchio, A. A. J. Hamers, C. C. Hedrick, Nonclassical monocytes in health and disease. Annual review of immunology 37, 439--456 (2019).

43. H. Zhu et al., CD16+ monocyte subset was enriched and functionally exacerbated in driving T-cell activation and B-cell response in systemic lupus erythematosus. Frontiers in immunology 7, 512 (2016).

44. A. Brunet et al., NR4A1-dependent Ly6Clow monocytes contribute to reducing joint inflammation in arthritic mice through Treg cells. European journal of immunology 46, 2789--2800 (2016).

45. V. Atsaves, V. Leventaki, G. Z. Rassidakis, F. X. Claret, AP-1 transcription factors as regulators of immune responses in cancer. Cancers 11, 1037 (2019).

46. G. J. Martinez, R. M. Pereira, ij, The transcription factor NFAT promotes exhaustion of activated CD8+ T cells. Immunity 42, 265--278 (2015).

47. F. Macian et al., Transcriptional mechanisms underlying lymphocyte tolerance. Cell 109, 719--731 (2002).

48. E. J. Wherry et al., Molecular signature of CD8+ T cell exhaustion during chronic viral infection. Immunity 27, 670--684 (2007).

49. G. Galletti et al., Two subsets of stem-like CD8+ memory T cell progenitors with distinct fate commitments in humans. Nature immunology 21, 1552--1562 (2020).

50. R. Wang et al., The transcription factor Myc controls metabolic reprogramming upon T lymphocyte activation. Immunity 35, 871--882 (2011).

51. W. Seo et al., Runx-mediated regulation of CCL5 via antagonizing two enhancers influences immune cell function and anti-tumor immunity. Nature communications 11, 1562 (2020).

52. G. I. Welsh, S. Miyamoto, N. T. Price, B. Safer, C. G. Proud, T-cell Activation Leads to Rapid Stimulation of Translation Initiation Factor eIF2B and Inactivation of Glycogen Synthase Kinase-3. Journal of Biological Chemistry 271, 11410--11413 (1996).

53. A. Taylor et al., Glycogen synthase kinase 3 inactivation drives T-bet-mediated downregulation of co-receptor PD-1 to enhance CD8+ cytolytic T cell responses. Immunity 44, 274--286 (2016).

54. H. Chi, Regulation and function of mTOR signalling in T cell fate decisions. Nature reviews immunology 12, 325--338 (2012).

55. S. S. Iyer, G. Cheng, Role of interleukin 10 transcriptional regulation in inflammation and autoimmune disease. Critical Reviews™ in Immunology 32, (2012).

56. H. Lv, C. Lai, W. Zhao, Y. Song, GABPB1-AS1 acts as a tumor suppressor and inhibits non-small cell lung cancer progression by targeting miRNA-566/F-box protein 47. Oncology Research 29, 401 (2021).

57. W. Qi et al., LncRNA GABPB1-AS1 and GABPB1 regulate oxidative stress during erastin-induced ferroptosis in HepG2 hepatocellular carcinoma cells. Scientific Reports 9, 16185 (2019).

58. S. Ren et al., RNA-seq analysis of prostate cancer in the Chinese population identifies recurrent gene fusions, cancer-associated long noncoding RNAs and aberrant alternative splicings. Cell research 22, 806-821 (2012).

59. C. Hao et al., Up-regulation of ANKDR49, a poor prognostic factor, regulates cell proliferation of gliomas. Bioscience Reports 37, BSR20170800 (2017).

60. Y.-h. Liu et al., ANKRD49 promotes the invasion and metastasis of lung adenocarcinoma via a P38/ATF-2 signalling pathway. Journal of Cellular and Molecular Medicine 26, 4401--4415 (2022).

61. B. Schoeps, J. Frdrich, A. Krger, Cut loose TIMP-1: An emerging cytokine in inflammation. Trends in cell biology 33, 413--426 (2023).

62. A. Adamson et al., Tissue inhibitor of metalloproteinase 1 is preferentially expressed in Th1 and Th17 T-helper cell subsets and is a direct STAT target gene. PLoS One 8, e59367 (2013).

63. J.-L. Hsieh et al., CD8+ T cell-induced expression of tissue inhibitor of metalloproteinses-1 exacerbated osteoarthritis. International Journal of Molecular Sciences 14, 19951--19970 (2013).

64. (Alliance of Genome Resources), vol. 2023.

65. Harmonizing model organism data in the Alliance of Genome Resources. Genetics 220, iyac022 (2022).

66. M. Houtman et al., T-cell transcriptomics from peripheral blood highlights differences between polymyositis and dermatomyositis patients. Arthritis research \& therapy 20, 1--15 (2018).

67. Y. Wu et al., Cytoskeleton regulator RNA expression on cancer-associated fibroblasts is associated with prognosis and immunotherapy response in bladder cancer. Heliyon 9, (2023).

68. Z. Li et al., Single-cell RNA-seq reveals characteristics of malignant cells and immune microenvironment in subcutaneous panniculitis-like T-cell lymphoma. Frontiers in Oncology 11, 611580 (2021).

69. X. Wang et al., The long non-coding RNA CYTOR drives colorectal cancer progression by interacting with NCL and Sam68. Molecular cancer 17, 1--16 (2018).

70. S. Kurella et al., Transcriptional modulation of TCR, Notch and Wnt signaling pathways in SEB-anergized CD4+ T cells. Genes \& Immunity 6, 596--608 (2005).

71. D. Elewaut, C. F. Ware, The unconventional role of LT$\alpha$$\beta$ in T cell differentiation. Trends in immunology 28, 169--175 (2007).

72. V. Upadhyay, Y.-X. Fu, Lymphotoxin signalling in immune homeostasis and the control of microorganisms. Nature Reviews Immunology 13, 270--279 (2013).

73. K. V. Korneev et al., TLR-signaling and proinflammatory cytokines as drivers of tumorigenesis. Cytokine 89, 127--135 (2017).

74. M. Stoeckius et al., Cell Hashing with barcoded antibodies enables multiplexing and doublet detection for single cell genomics. Genome Biology 19, 224 (2018).

75. C. Morse et al., Proliferating SPP1/MERTK-expressing macrophages in idiopathic pulmonary fibrosis. European Respiratory Journal 54, (2019).

76. E. Valenzi et al., Single-cell analysis reveals fibroblast heterogeneity and myofibroblasts in systemic sclerosis-associated interstitial lung disease. Annals of the rheumatic diseases 78, 1379--1387 (2019).

77. S. Grebinoski et al., Autoreactive CD8+ T cells are restrained by an exhaustion-like program that is maintained by LAG3. Nature immunology 23, 868--877 (2022).

78. G. X. Y. Zheng et al., Massively parallel digital transcriptional profiling of single cells. Nature communications 8, 14049 (2017).

79. L. Waltman, N. J. Van Eck, A smart local moving algorithm for large-scale modularity-based community detection. The European physical journal B 86, 1--14 (2013).

80. C. Trapnell et al., The dynamics and regulators of cell fate decisions are revealed by pseudotemporal ordering of single cells. Nature biotechnology 32, 381--386 (2014).

81. X. Qiu et al., Reversed graph embedding resolves complex single-cell trajectories. Nature methods 14, 979--982 (2017).

82. J. Cao et al., The single-cell transcriptional landscape of mammalian organogenesis. Nature 566, 496--502 (2019).

83. V. Bergen, M. Lange, S. Peidli, F. A. Wolf, F. J. Theis, Generalizing RNA velocity to transient cell states through dynamical modeling. Nature biotechnology 38, 1408--1414 (2020).

84. V. A. Traag, L. Waltman, N. J. Van Eck, From Louvain to Leiden: guaranteeing well-connected communities. Scientific reports 9, 5233 (2019).

85. J. H. Levine et al., Data-driven phenotypic dissection of AML reveals progenitor-like cells that correlate with prognosis. Cell 162, 184--197 (2015).

86. C. Hafemeister, R. Satija, Normalization and variance stabilization of single-cell RNA-seq data using regularized negative binomial regression. Genome biology 20, 296 (2019).

87. R. Xiang et al., A comparison for dimensionality reduction methods of single-cell RNA-seq data. Frontiers in genetics 12, 646936 (2021).

88. A. Gayoso et al., A Python library for probabilistic analysis of single-cell omics data. Nature biotechnology 40, 163--166 (2022).

89. C. Xu et al., Probabilistic harmonization and annotation of single-cell transcriptomics data with deep generative models. Molecular systems biology 17, e9620 (2021).

90. M. D. Luecken et al., Benchmarking atlas-level data integration in single-cell genomics. Nature methods 19, 41--50 (2022).

91. M. Hu, M. Chikina, InstaPrism: an R package for fast implementation of BayesPrism. bioRxiv, 2023--2003 (2023).

